# Metagenomic dynamics in *Olea europaea* after root damage and *Verticillium dahliae* infection

**DOI:** 10.1101/555185

**Authors:** Jose Manuel Martí, Luis F. Arias, Wladimiro Díaz, Vicente Arnau, Antonio Rodriguez-Franco, Carlos P. Garay

## Abstract

The olive tree is of particular economic interest in the Mediterranean basin. Researchers have conducted several studies on one of the most devastating disorders affecting this tree, the Verticillium wilt of olive, which causes significant economic damage in numerous areas of this crop. We have analyzed the temporal metagenomic samples of a transcriptomic study in *Olea europaea* roots and leaves after root-damage and after a root *Verticillium dahliae* infection (Jimenez-Ruiz et al. 2017). Our results indicate that this infection, although led by *Verticillium*, is driven not by a single species but by a polymicrobial community, including their natural endophytes, which acts as a consortium in the attack to the host plant. This community includes both biotrophic and necrotrophic organisms that alternate and live together during the infection. Our results not only describe how the microbial community progresses along these processes, but also explain the high complexity of these systems, that in turn, could justify at least in part the occasional changes and disparity found at the time of classifying the kind of parasitism of a determined organism.

## Introduction

The olive tree could be the earliest cultivated temperate fruit since paleobotanists have traced back its domestication to the early Neolithic age (Terral and Arnold-Simard 1996). At present, both the cultivation of olive and the olive-oil related industry have grown to the point of having a profound worldwide socioeconomic and environmental impact.

As the Verticillium wilt of olive is one of the most devastating disorders affecting this crop, a transcriptomic RNA-seq analysis was recently conducted to study the interaction between *Olea europaea* and *Verticillium dahliae* (Jimenez-Ruiz et al. 2017), which concluded that mainly a ROS response appeared first in the pathogen and later in the plant.

In recent years, there has been a deviation from studying individual species to study the whole community that is actually living in a determined niche. Shotgun metagenomic sequencing (SMS) helps this paradigm shift by the use of massive nucleic acid sequencing of whole metagenomes obtained from samples with the extra advantage of not requiring prior knowledge of the species that are present.

Soil microorganisms constitute one of the most complex systems in nature, where many different life forms interact one to another. Fungi play a major role as they act as parasites, saprotrophs or mutualists in a myriad of environments, including the rhizosphere of many different plants (Cuadros-Orellana et al. 2013). The description of these interactions are, however, variable and somehow confusing in many cases. So, *V. dahliae* has been described as an hemi-biotrophic pathogen because it seems to behave as biotrophic during the initial stages of plant infection, but changes to a necrotrophic lifestyle during subsequent stages (Scholz et al. 2018; Shaban et al. 2018; Häffner and Diederichsen 2016). The same goes for other fungus like those pertaining to the *Fusarium* gender (Lyons et al. 2015; Yang et al. 2013). However, the molecular or biological bases underlying these different parasitical alternations are still not fully understood.

We must emphasize that the use of metagenomic data dealing with the process of a fungal plant infection is almost lacking. A search in the ENA metagenomic database *(Mgnify* database) for fungal infections in plants shows only one hit (case `MGYS00001376`) that deals with the still unpublished study of the infection by *Erysiphe alphitoides*, the causal agent of oak powdery mildew, and other foliar microorganisms of pedunculate oak *(Quercus robur* L.). Some recent studies (Donovan et al. 2018) focus on the characterization of fungal metagenomics in animal species, especially those from pig and mouse microbiomes. Human metagenomics, particularly that related to gut microbiome, has experienced a recent burst, mainly by the unexpected conse quences for health and disease (Gilbert et al. 2018), which are especially evident through time series analysis (Martí et al. 2017). However, generally speaking, animal, plant and fungal metagenomes are not sufficiently studied in comparison with bacterial microbiomes.

We have reanalyzed the samples obtained by Jimenez-Ruiz et al. (2017) with the new perspective of longitudinal (temporal) metagenomics to unravel the dynamics of the infection. Our results indicate that the Verticillium wilt of olive is a complex infection process involving more contenders than just *Olea* and *V. dahliae.* We show that along the infection under natural conditions, there is a biological succession of different kind of parasites (mainly biotrophic and necrotrophic) that could at least partially explain the observed parasitic alternations described in many systems. A careful design of the experimental conditions, such as the using of sterile seeds and substrates to raise the plants ensuring that only the parasite is growing, is needed to be able to reach the correct conclusions.

## Methods

The complete RNA-seq dataset of the study by Jimenez-Ruiz et al. (2017) consisted in a 2×100 paired-ends and unstranded library that was downloaded from the NCBI SRA servers with accession numbers provided in the article (Jimenez-Ruiz et al. 2017).

### Pre-analysis

RNA-seq SMS data was quality checked using FastQC v0.11.5 (Babraham Bioinformatics 2018) and MultiQC v1.3 (Ewels et al. 2016).

### Mapping of reads

The whole SMS library was mapped against the olive and the *Verticillium* genomes per separate using both the pseudomapper Kallisto v0.44 (Bray et al. 2016) and the RNA-seq aligner STAR v2.7 (Dobin et al. 2013).

### SMS database preparation

The database used for the Centrifuge program was generated in-house from the complete NCBI `nt` database (nucleotide sequence database, with entries from all traditional divisions of GenBank, EMBL, and DDBJ) and index databases (Wheeler et al. 2007), downloaded in Dec 2017. This database was supplemented with all the sequences in the NCBI whole genome shotgun `WGS` database (Wheeler et al. 2007) belonging to the *Olea* genus and the fungi kingdom. Once generated, the Centrifuge indexed compressed database weighted more than 135 GB. So far, this is the most massive Centrifuge database we have prepared and used successfully.

### Classification

The metagenomic sequences were analyzed with the Centrifuge software package (Kim et al. 2016) version 1.0.3-beta (Dec 2017), run in parallel within a shared-memory fat node, using 8 threads and peaking half a tebibyte of DRAM.

### Post-analysis

The results generated with Centrifuge were post-processed, analyzed and visualized using `Recentrifuge` (Marti 2018), release v0.22.1 (Oct 2018). This parallel software was run with the flags -- **minscore 50** (`mhl` set to 50) and -**x CMPLXCRUNCHER** to prepare the `Recentrifuge` quantitative output to be introduced in `cmplxCruncher` *(in preparation)* as input. A recent development version of `cmplxCruncher` (Oct 2018) was used to perform the final temporal metagenomic analysis and produce the plots shown.

## Results and Discussion

### Quality check and mapping of the SMS reads

The analysis with FastQC and MultiQC showed a notorious change in the per base GC content in the infected root RNA samples that increased with time after the inoculation (Figure 1a). The GC content in the sequenced reads evolved from a unimodal distribution peaking at 43%, coincident with that of the olive genome, to a pseudogaussian distribution reflecting a higher GC content ratio (53%) 15-day after the inoculation. Figures 1b and 1c show the percentage of reads coming from infected roots mapped either with Kallisto or STAR to the olive genome drastically decreasing along the infection. The number of *Verticillium* reads was very low in the infected samples. As expected, *Verticillium* reads were negligible or missing in the controls and leaves. However, the per base GC content and the proportion of mapped reads in the leaf samples to the olive genome remained practically constant. All these data taken together confirmed that the infection progressed through the roots and that there was a progressive emergence of other biological organisms during infection that were displacing the olive tree in terms of mRNA abundance. The species origin of the unmapped reads was unknown, and thus, a metagenomic analysis was required.

**Figure 1.**
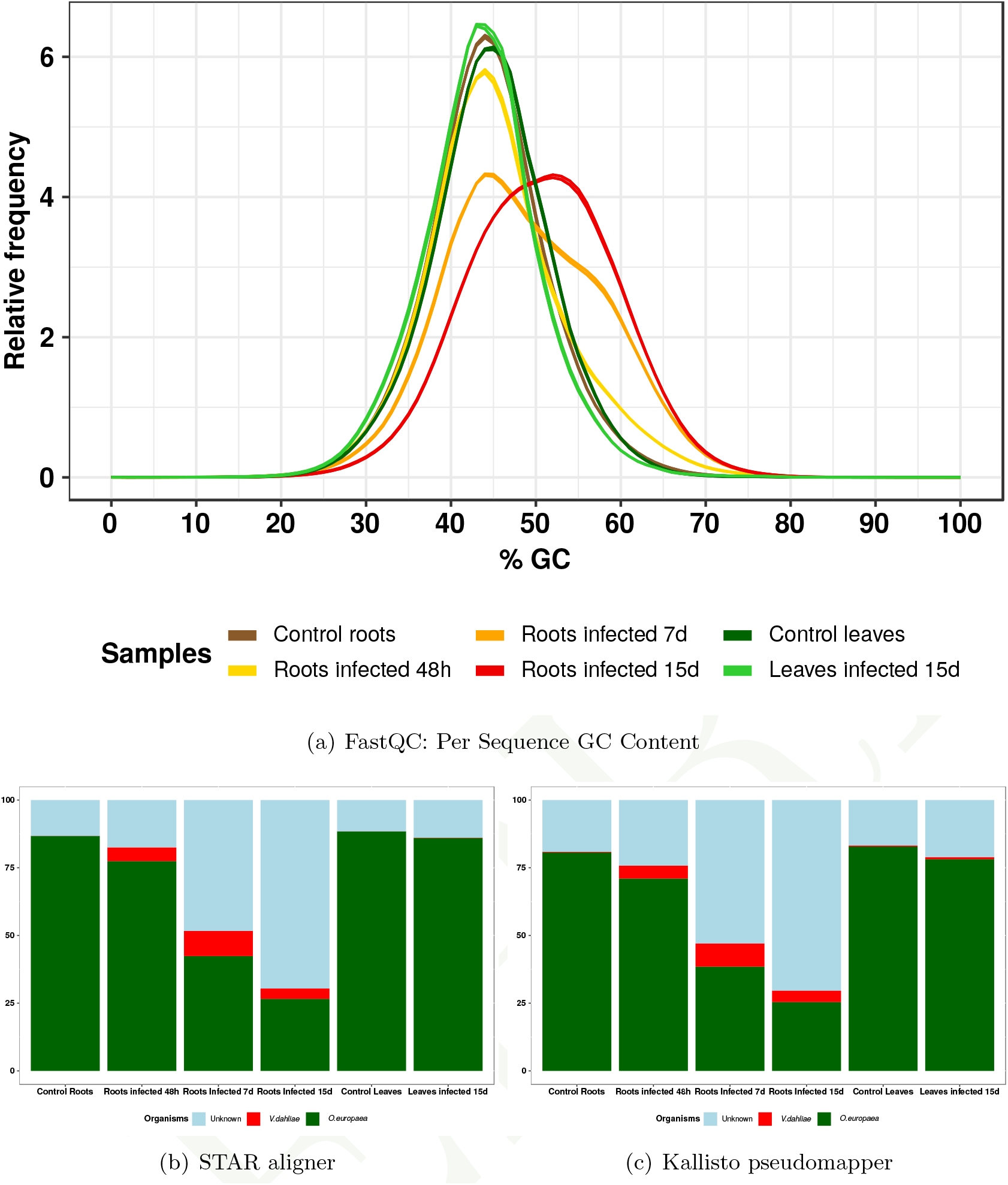
(a): The average per base GC content of the reads. The GC content of the olive genome is roughly 43%. (b-c): Percentage of reads mapped to the olive (green) and *V.dahliae* (red) genomes, obtained through STAR and Kallisto, respectively. The cyan color corresponds to the proportion of unmapped reads of initially unknown origin.

### Temporal metagenomic analysis of the root infection process Overall dynamics of the infection

Figure 2 shows how the infection with *V. dahliae* caused a profound impact in the rhizosphere of the olive roots. This is shown by comparing the rank dynamics of mapped reads and the stability plot for species of the roots control sample (first column) to the roots 48 hours after the infection (second column). The boost in the relative frequency of *V. dahliae* reads is the most obvious, but it is not the only change. The same figure shows that other species are taking advantage of the *Verticillium* rise, and some others are suffering an apparent displacement. The low values in the rank stability index —RSI— (Martí et al. 2017) column and the extreme fluctuations in the RSI plot indicate that the rhizosphere experienced an intense perturbation with the inoculation of *V. dahliae*, also corroborated by peak values both in rank and differences variability plots. Thereafter, the rhizosphere was undergoing a transient state as a complex system. The instability is reflected in the lower taxa present in all samples compared to the non-perdurable taxa along the infection (see Supplemental Figure 10).

**Figure 2.**
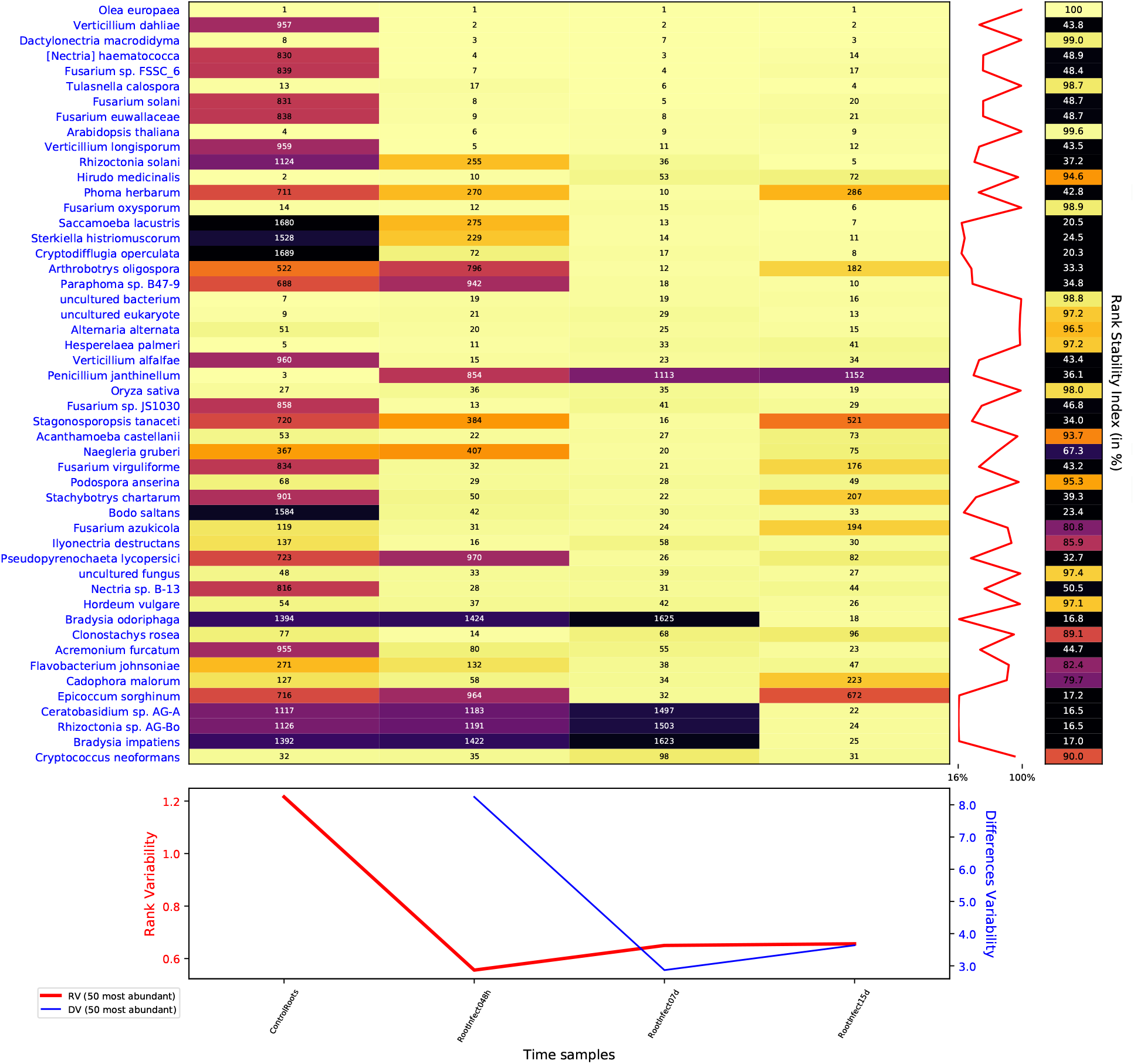
Rank dynamics and stability plot for mapped reads classified at species level during the process of infection with *V. dahliae.* The dynamics of the rank during the process shows the profound impact in the rhizosphere of the olive tree caused by the inoculation with *V. dahliae.* Numbers and colours (using perceptually uniform colormap for easier visualization) show the ranking by the accumulated species abundance in each column. Different rank variability and stability measurements (Martí et al. 2017) are given. The right panel shows the rank stability throughout species ordered by their overall abundance. The lower panel contains plots of the rank variability along time.

Figure 3, the clustered correlation and dendrogram plot for mapped reads at species level during the infection, shows the evident antagonism between the cluster formed by *Olea europaea* and *Penicillium janthinellum*, and the cluster formed by *Verticillium* spp., located in opposite extremes of the clustered correlation matrix. As indicated in the figure, the clusterization algorithm formed five superclusters with a total of 20 clusters.

**Figure 3.**
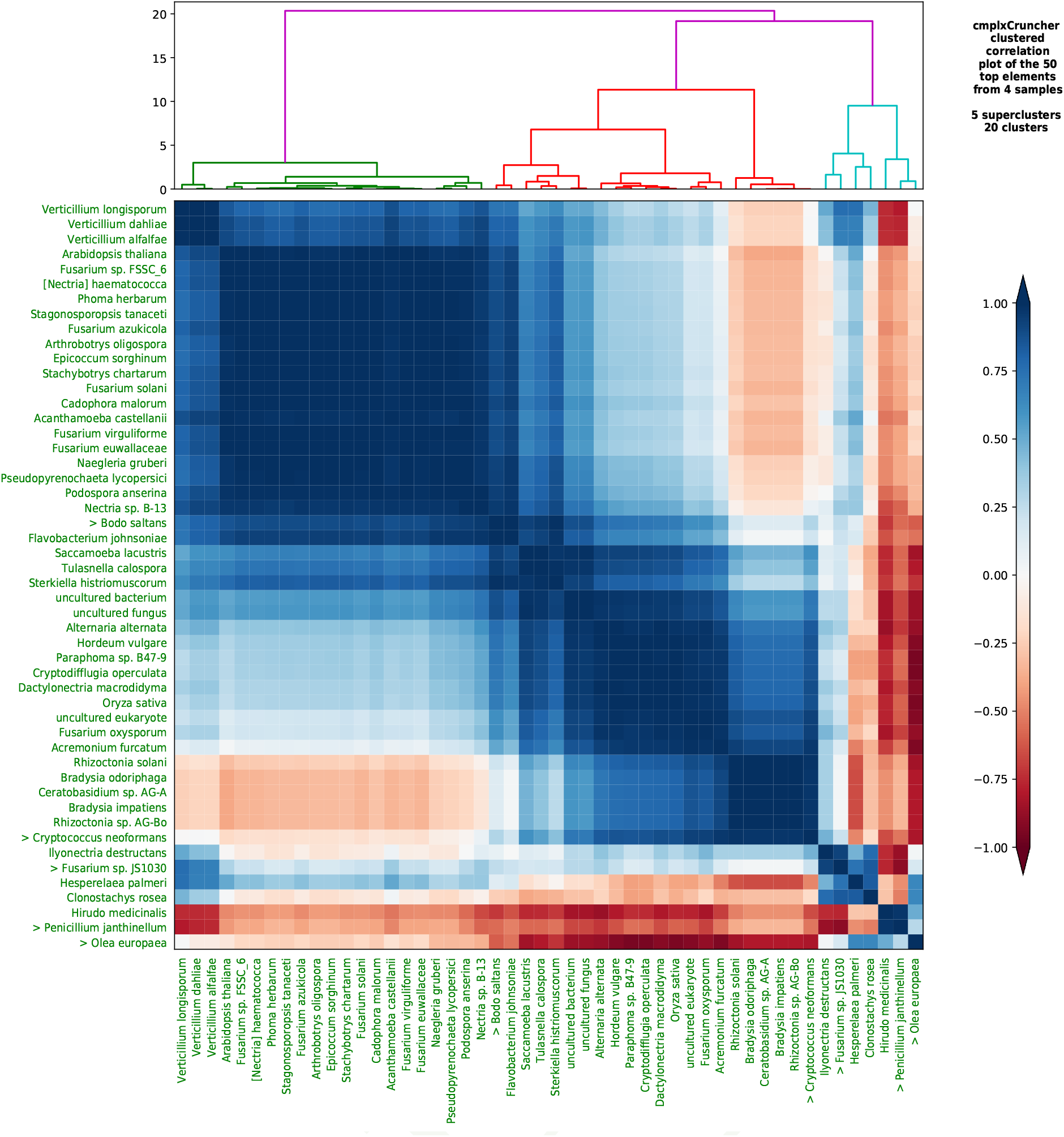
Clustered correlation and dendrogram plot for species during the process of infection with *V. dahliae.* We show the 50 most abundant species ordered by clusterization based on the Pearson time correlation matrix. Several clusters and superclusters can be identified with this analysis by `cmplxCruncher`.

Based on our methods for the analysis of microbiota variability and stability (Martí et al. 2017), we have fit a power-law to std *σ_i_* vs. mean *μ_i_* for the relative abundance of genera during the root infectious process (see Figure 4). The scaling index *β* ~ 1 of this Taylor’s law indicates that the biological system follows the model of an exponential or a geometric distribution, which are characterized by *β* = 1. In addition, Supplemental Figure 14 plots Taylor’s law parameter space with data from the fits for different taxonomic ranks performed for the dataset of roots olive infection with *V. dahliae.* We can see that there is a correlation between *β* and *V* depending on the taxonomic level. We can see how the `no_rank` subsample, with no separation by taxonomic level, is located in an intermediate position.

**Figure 4.**
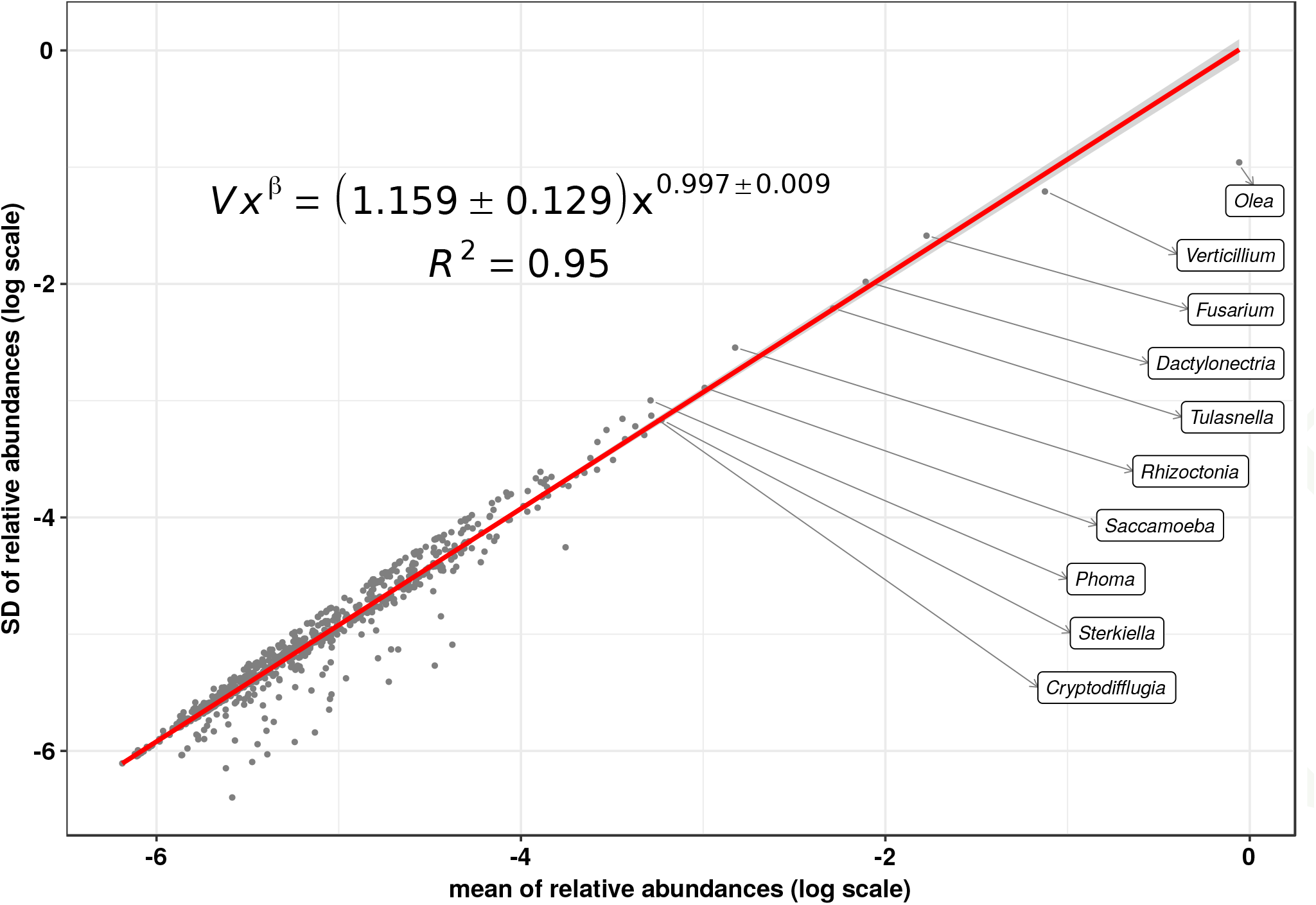
Taylor’s law of the biological system consisting in the metagenome at genus level along the root infectious process. We see that Taylor’s power law spans six orders of magnitude, therefore, it is ubiquitous.

The `GeneFiltløk` subsample is an experimental dataset containing the 10,000 most representative expressed-genes in each sampling time. Supplemental Figure 15 depicts the x-Weighted fit in logarithmic scale (see Martí et al. (2017) for details on the fit), which shows a modest generalized coefficient of determination 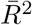 because this dataset has an important component of variability unaccountable by the power-law model. This dispersion of the residuals is evident in Supplemental Figure 16, the residual analysis plot of the correspondent LLR-model fit, which in addition shows that the residuals are normally distributed but present some degree of deviation from homoscedasticity. Finally, Supplemental Figure 12 shows the `Recentrifuge` plots of fungal SMS classified reads at species level for leaves of two specimens, the control sample and the sample 15 days after the *Verticilium* inoculation.

In the following subsections, we show the evolution during the infection for some significative clades: Amoebae and Cilliates, Fungi, Bacteria, and Nematoda.

### Amoebae and Ciliates

The *Verticillium* infection likely broke trophic network equilibria because direct or indirectly could cause breakage, destructuring of tissues and lysis of cells, thus promoting the grown of opportunistic organisms. In this respect, in Figure 2 we can see how three protist species *(Saccamoeba lacustris, Sterkiella histriomuscorum*, and *Cryptodifflugia operculata)*, which were not among the 1500 most frequent species in the roots control sample, evolve during the infection to be among the 15 most frequent species found by number of SMS reads assigned. In the rhizosphere, the ubiquitous free-living *Saccamoebae* are living in biofilms and at the interfaces between roots and water (Kebbi-Beghdadi and Greub 2014). The ciliate *Sterkiella histriomuscorum* (before known as *Oxytricha trifallax)* is a cosmopolitan species in soil, but it is also habitual in limnetic habitats (Foissner and Berger 1999; Kumar et al. 2017). The amoeba *Cryptodifflugia operculata* is a bacterivore which is also able to prey on larger nematodes thanks to efficient and specialized cooperative hunting (Geisen et al. 2015).

Certain amoeboid protists are pathogenic for the olive. Concretely, some slime molds of the genus *Didymium* are associated with a severe disease of the olive flower-buds, causing extensive destruction and blockage of flower development (Medeira et al. 2006). During the infection process, *Didymium* spp. (particularly *D. squamulosum* and *D. iridis)* appeared on day 7, and remained on day 15. In fact, reads belonging to the Myxogastria class, which contains the genus *Didymium*, increased 3.8 times from the control sample to 48h after infection, but they rocketed 10.8 times from 48 h to 7 days after the infection. Taking into account the variation in the absolute number of reads assigned for each sample (see Supplemental Figure 11), Myxogastria relative frequency is 2 × 10’^6^ in the control and about 1 × 10’^4^ seven days after inoculation. Such a growth is probably due to the increased availability of decaying plant material as a consequence of the appearance of destructive necrotrophic species that took advantage of the *V. dahliae* isolate V937I inoculation, which is an archetype of the highly virulent D pathotype (Jimenez-Ruiz et al. 2017).

### Fungi

Figure 5 is a collection of four `Recentrifuge` plots (Marti 2018) showing the evolution of fungal SMS reads during the *V. dahliae* infection of the olive roots. *Penicillium janthinellum* dominates the roots control sample before the infection. *P. janthinellum* is an endophytic fungus which seems to be remedial to plants in the alleviation of heavy metal stress by enhancing the host physiological status. Therefore, it is not by chance that *Olea europaea* and *Penicillium janthinellum* appear clustered together in Figure 3.

As expected, the frequency of mapped reads indicated that *V. dahliae* became the dominant fungus in the roots soon after the inoculation. Its number of reads were the second most frequent just after those pertaining to the olive host (see Figure 2 and Figure 5). Nevertheless, in the following samples, without losing the second position, its relative frequency started to decrease in favor of other fungi (see Figure 5) that we review below.

**Figure 5.**
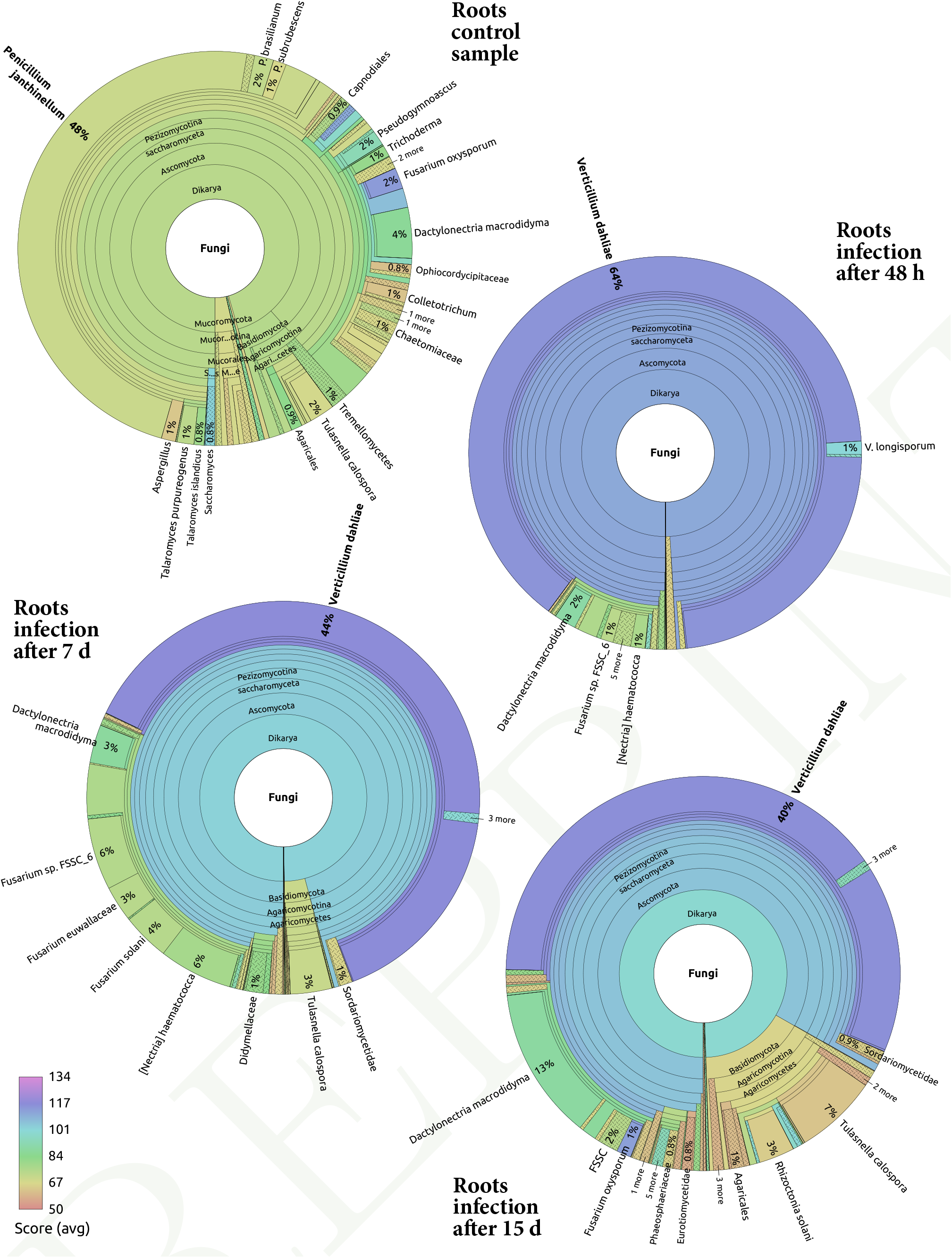
Recentrifuge plots of the evolution of fungal SMS classified reads at species level during the *V. dahliae* infection. The top pie corresponds to the root control sample, the others to the infected roots after 48 h, 7 days, and 15 days, respectively, of the *Verticilium* inoculation. An interactive and dynamic collection of `Recentrifuge` plots (Marti 2018) can be accessed via the official project’s webpage at https://www.uv.es/marti?m/olea.

*Dactylonectria macrodidyma* is a fungus that was already present in the control sample, but which benefited from the *V. dahliae* infection since it becomes the third most frequent species in the last temporal point (see Figure 5) just behind the host and the inoculated fungi (see Figure 2). *D. macrodidyma* is itself another pathogenic and necrotrophic fungus in crops as it is the causative agent of root rot disease of many herbaceous and woody plants such as grapevine, avocado, cherimola and olive (Malapi-Wight et al. 2015), some of them fruit trees whose top exploiters are Spain and Chile (Auger et al. 2015). *Tulasnella calospora* is another similar case to *D. macrodidyma* since it was present in the control sample but ended up as the fourth most frequent species in the time series of the infection. *T. calospora* has been recently studied as a mycorrhizal fungal symbiont of orchids (Fochi et al. 2017), but in our case, it seemed somehow to take advantage of the *V. dahliae* infection probably due to the destructuration, destruction or lysis of tissues and cells. In fact, species of Tulasnellaceae have been described to be both symbionts and saprotrophs simultaneously (Sarmiento 2016).

One week after the inoculation, reads assigned to the so named *Fusarium solani* species complex (FSSC) represent a fifth of all the fungal reads (see Figure 5). *Nectria haematococca* and its asexual counterpart, *Fusarium solani*, are the most relevant species in this complex. While researchers in Spain have reported that *F. solani* is only weakly pathogenic on olive (Hernández et al. 1998), this fungus has caused fatal wilt of *Olea europaea* in Nepal (Vettraino et al. 2009).

Additionally, the infected samples also contain *Fusarium euwallaceae*, a genealogically exclusive lineage of fungi within Clade 3 of the FSSC discovered as a fungal symbiont of *Euwallacea* sp., an invasive ambrosia beetle that causes serious damage to more than 20 species of olive tree (Freeman et al. 2013). This taxon, with average score below the paired-ended read half value (100) may represent another close species in the FSSC.

Another frequent *Fusarium* fungi in the studied samples, *Fusarium oxysporum*, is the causal agent of the Fusarium wilt in many different plants, including tomato, chickpea, and others (García-Pedrajas et al. 1999), but it is considered only slightly pathogenic for olive in Spain (Hernández et al. 1998). In fact, it is present in the control root samples and keeps a similar rank throughout infection process, except for the last to 15 days, where it has advanced to the sixth position of all species (see Figure 2). Generally speaking, *Fusarium oxysporum* is one of those cases in which a debate is open as to whether this fungus is considered a biotroph, a hemibiotroph, or a necrotroph, able to kill plant tissue quickly and thereafter feeding saprotrophically on the dead remains (Edgar et al. 2006; Thaler et al. 2004; Ma et al. 2013; Moore et al. 2011).

The fungi *Rhizoctonia solani*, R. sp. AG-Bo, and *Ceratobasidium* sp. AG-A belongs to the same cluster (see Figure 3). As we can see in Figure 2, these fungi presented a very low frequency of mapped reads during the time series except in the last sample corresponding to 15 days after the inoculation with *V. dahliae. Rhizoctonia solani* is a soil-borne plant pathogen that has been related to rotten roots in olives (Hernández et al. 1998). Both *Rhizoctonia* and *Ceratobasidium* genera belong to the family Ceratobasidiaceae of saprotrophic and cosmopolitan fungi which could be facultative plant pathogens with a wide host range (Moore et al. 2011).

### Bacteria

The bacterial content of the samples was severely shrunk because the mRNA was isolated using poly(A) columns (Jimenez-Ruiz et al. 2017), and it is hence, biased. Despite that limitation, the overall dynamics of the bacterial community may still be outlined for the infection. Figure 6 shows the rank dynamics and stability plot for bacterial species. The main difference with Figure 2, the overall rank dynamics and stability plot for species, dominated by fungi, is the position of the differences variability —DV— (Martí et al. 2017) peak. In the latter case, it is on the second sample (48 hours after the infection), while in the former case it appears over the third sample (one week after the inoculation). That means that the effects of the infection reached the bacterial community with some delay compared to the whole species population. The fact that the minimum in DV is on the third sampling time for the whole population but on the fourth sampling time for bacteria supports the existence of such delay.

**Figure 6.**
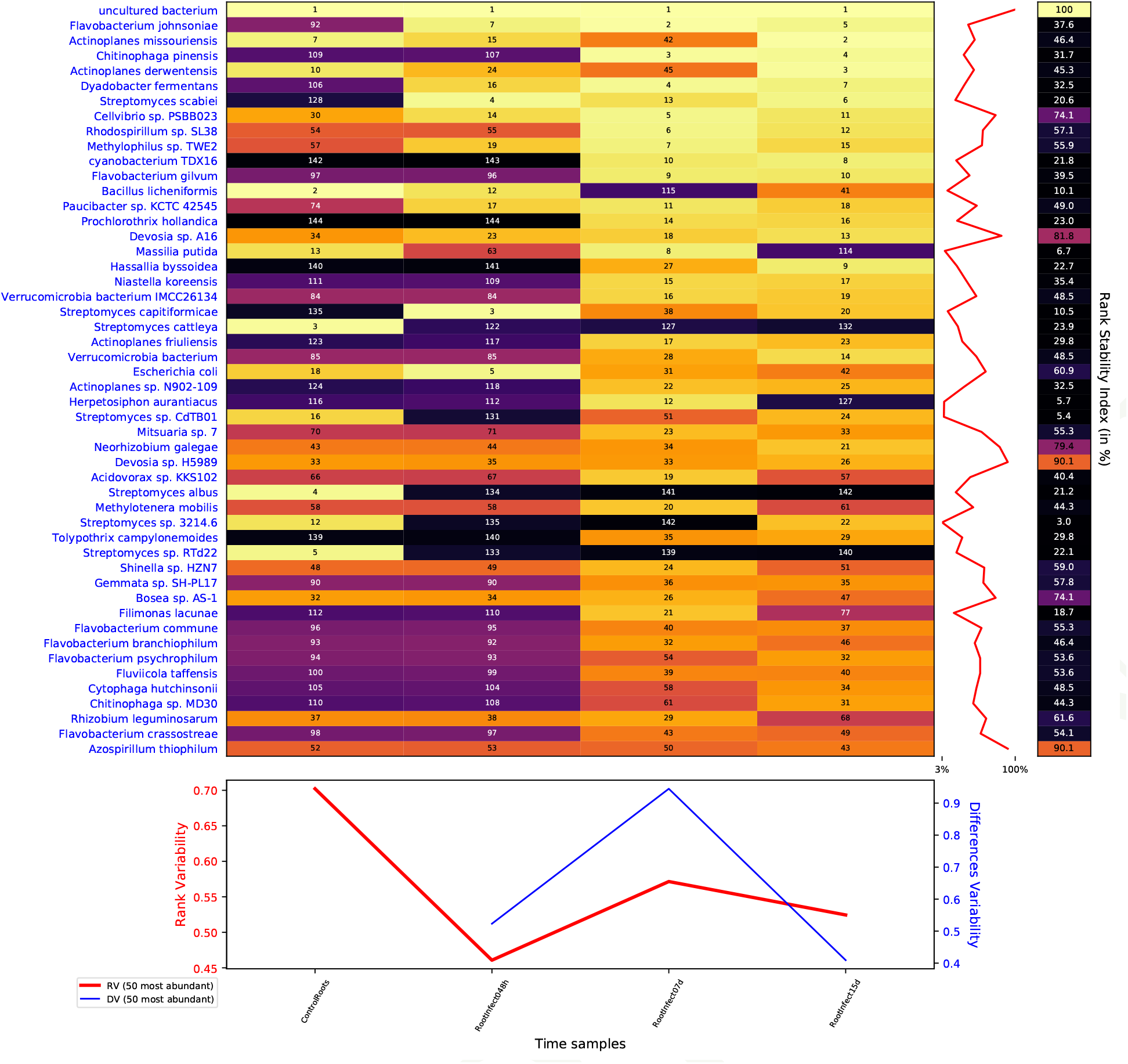
Rank dynamics and stability plot for bacterial species during the process of infection with *V. dahliae*. Numbers and colours (using perceptually uniform colormap for easier visualization) show the ranking by the accumulated species abundance in each column. Different rank variability and stability measurements (Martí et al. 2017) are given. The right panel shows the rank stability throughout species ordered by their overall abundance. The lower panel contains plots of the rank variability along time.

In Figure 6, the RSI shows low values compatible with the perturbation introduced in the bacterial community with the inoculation of *V. dahliae.* Intriguingly, a couple of *Devosia* species (sp. A16 and sp. H5989) are exceptions to this behavior since present high RSI of 90% and 82%, respectively.

Other entirely different cases are *Chitinophaga pinensis* and *Flavobacterium johnsoniae*, which were not very common in the first two samples but then moved forward more than 100 rank positions to reach the top 4 and top 5 in the last two sampling times, respectively. Both are soil-borne bacteria that belong to the widespread and diverse Bacteroidetes phylum and are recognized for its capacity to degrade chitin, the main component in the exoskeleton of arthropods and the cell walls of fungi, so that they could be endohyphal bacteria (a kind of endosymbiont) of fungi belonging to the *F. solani* species complex (Shaffer et al. 2017). Another relevant possibility is that those bacteria could have been recruited by the olive tree through root exudates as an indirect plant defense mechanism against the fungal attack (Baetz 2016; Häffner and Diederich-sen 2016). Indeed, chitinolytic bacteria are well-known antagonists of plant pathogenic fungi (Frändberg and Schnürer 1998). In fact, the rhizosphere of wild olive is a reservoir of bacterial antagonists of *V. dahliae* showing chitinolytic activity (Aranda et al. 2011). The dynamics of *C. pinensis* and *F. johnsoniae* shown in Figure 6 and the dynamics of species belonging to the FSSC demonstrated in Figure 2 seem compatible with such hypothesis.

### Nematoda

About Nematoda, it is remarkable the detection of *Oscheius tipulae* both with a high score and relatively high abundance on the sample of infected root after seven days. It also appears in the specimen of 8 h after root damage and, with lower abundance, 15 days after the infection. *O. tipulae* is one of the most common and cosmopolitan nematode species in soil (Félix 2006). While there is no clear relationship between this nematode and the infection dynamics in this study, it is also true that plants under attack are favoured by soilborne mobile predators such as nematodes, which are efficiently attracted by root-emitted compounds (Baetz 2016)

With low frequency and modest score, but with more biological significance, it was the finding of *Heterodera* exclusively in samples corresponding to 7 and 15 days after the inoculation into roots with *Verticillium* (see Supplemental Figure 13). *Heretodera* spp. are well-known plant-parasitic nematodes (PPN) associated with the olive tree, especially in nurseries, with reported cases in Spain (Ali et al. 2014). Other PPN such as *Meloidogynidae incognita* and *Pratylenchidae vulnus* (not found in the samples of this study) have been associated with *V. dahliae* synergistic coinfections to olives since it seems that, with the indirect root damages that they inflict on the trees, these nematodes act as the spearhead of other pathogenic soil-borne microor ganisms like *Verticillium.* Interestingly, Castillo et al. (1999) suggested that *Heretodera* and *Verticillium* may cooperate synergistically in the Verticillium wilt infection to yield both more widespread and more serious harm to the crop (Castillo et al. 1999). Our results point precisely in that direction. Finally, with low score, *Bursaphelenchus* also appears in Supplemental Figure 13. *Bursaphelenchus* spp. feeds on cells of trees by using stylets that pierce cell walls thanks to industrially useful *β*-glucosidases degrading enzymes, causing pests in palms and trees (Zhang et al. 2017).

### Temporal metagenomic analysis of the root damage process

Figure 7, the rank dynamics and stability plot for species, shows that the damage of the roots had a substantial effect on the rhizospheric microbiota but not as important as in the case of the infection with *V. dahliae* above. Comparing with Figure 2, we can see that the rank variability and especially the differences variability had lower values with the root damage than with the root infection.

**Figure 7.**
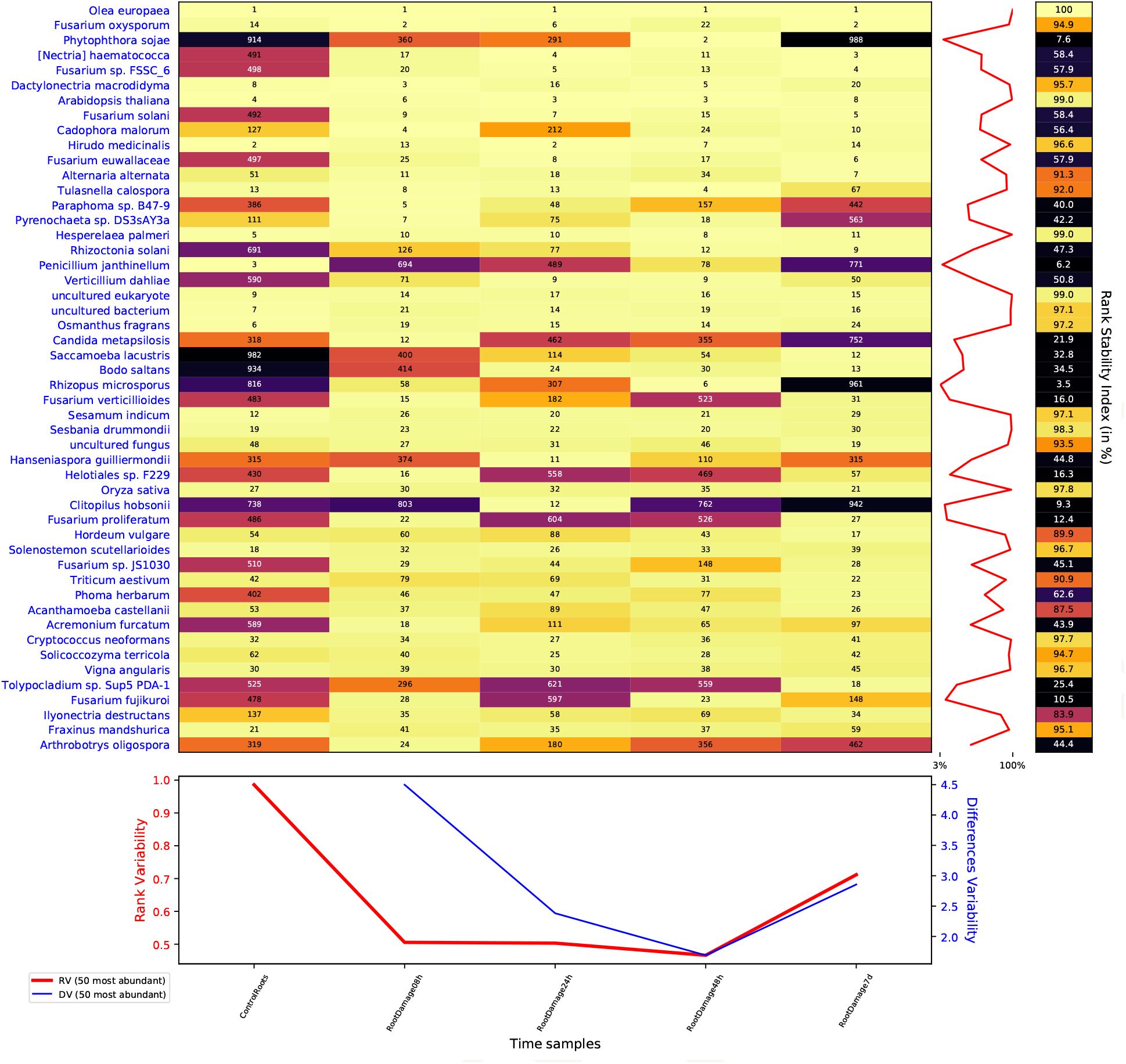
Rank dynamics and stability plot for species during the process after roots damage. The dynamics of the rank during the process shows a significative impact in the rhizosphere of the olive. Numbers and colours (using perceptually uniform colormap for easier visualization) show the ranking by the accumulated species abundance in each column. Different rank variability and stability measurements (Martí et al. 2017) are given. The right panel shows the rank stability throughout species ordered by their overall abundance. The lower panel contains plots of the rank variability along time.

However, there were also similarities in the evolution of both datasets despite their different timing. The dynamics of fungi belonging to the FSSC and the drop of *Penicillium janthinellum* abundance after damage are good examples. *P. janthinellum* abundance fell but the slump, being significant, was not as severe as in the infection. Fusarium spp. also benefit from the perturbation to the roots, growing as in the infectious case. *Verticillium dahliae* follows this same behavior even when it is in a different correlation cluster than FSSC species, as shows Supplemental Figure 17, the clustered correlation and dendrogram plot for species during the process after the roots damage.

Nevertheless, other taxa, at the end of the process (7 days), recovered a rank similar to the initial one. That is the case of the plant pathogen *Phytophthora sojae*, which causes root rot of soybean. *P. sojae* had a rank beyond 900 in the control samples, but it reached the second most frequent rank 48 hours after damage and returned to positions close to 1,000 in the last sample. *Rhizopus microsporus, Clitopilus hobsonii, Hanseniaspora guilliermondii*, and *Arthrobotrys oligospora* behaved similarly, having a final rank close to the initial one after a transient period. In particular, *H. guilliermondii* recovered precisely the same rank at the end (315).

Figure 8 shows the Taylor’s law fit for the relative abundance of genera throughout the root damage process. Comparing with Figure 4, we see a lower scaling index β and, interestingly, a much lower variability V. From a system dynamics perspective (Martí et al. 2017), these values indicate that the system is more stable after the root damage than after the inoculation with *Verticilium*, thus corroborating the above rank stability results.

**Figure 8.**
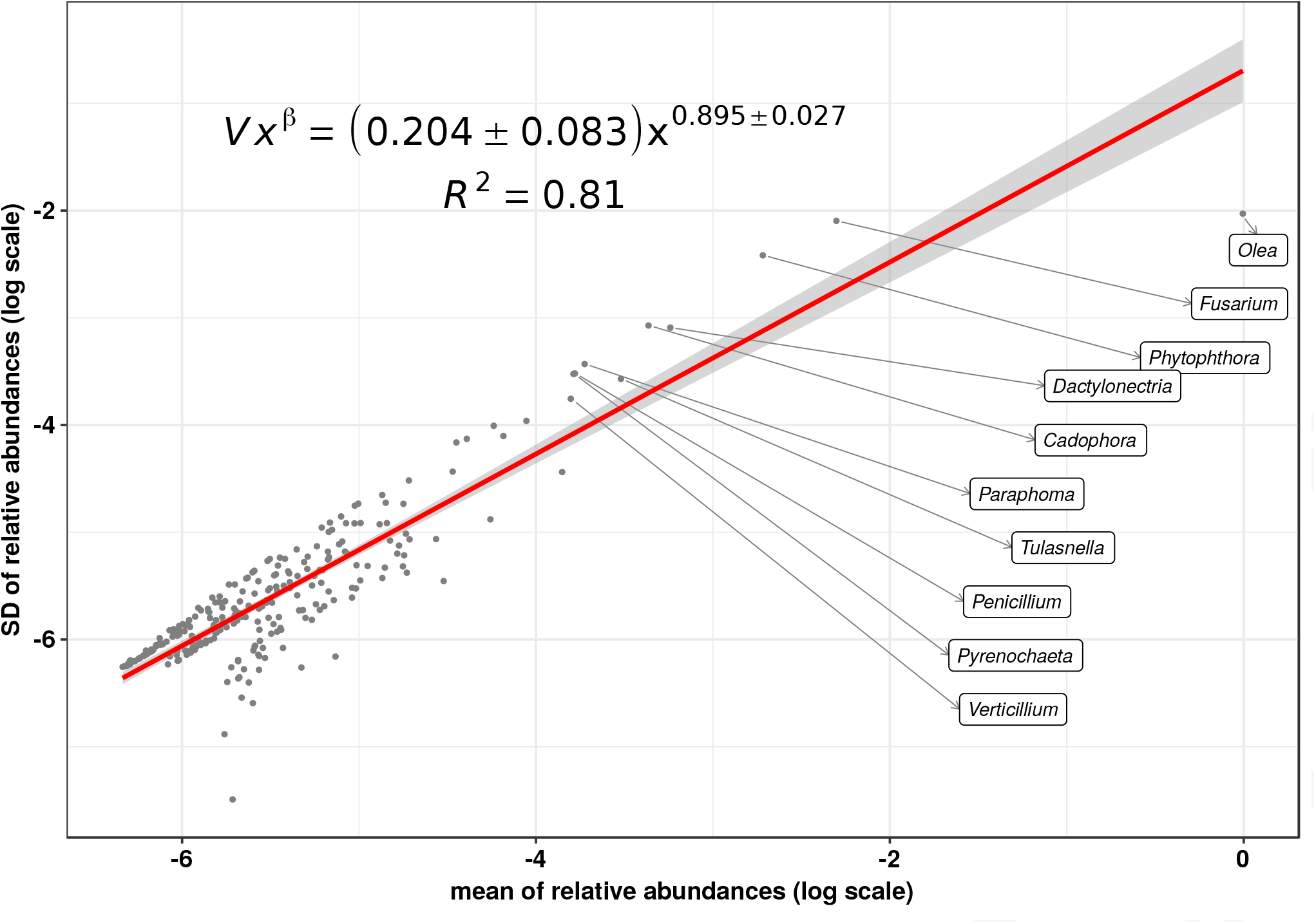
Taylor’s law of the biological system consisting in the metagenome at genus level throughout the root damage process. We see that Taylor’s power law seems to be ubiquitous, spanning in this case more than six orders of magnitude.

Finally, Supplemental Figure 18 shows the `Recentrifuge` plot of SMS classified reads for Dykaria fungi for the sample of leaves 15 days after the roots damage. *Candida albicans*, a known human pathogen that has been recently associated with ancient oaks too (Bensasson et al. 2019), appears with low frequency but good average confidence.

## Conclusions

Our results suggest that this disease, although led by *Verticillium*, is driven not by a single species but by a polymicrobial community which acts as a consortium in the attack of another community formed by the host plant and its endophytes, as Figure 9 shows. Thus, an infectious process can be generalized with a systems biology approach as an attack of one system, i.e. a polymicrobial community, to another, the host and its symbionts. Indeed, our systems approach to the Verticillium wilt of olive has revealed a complex interaction between complex systems.

**Figure 9.**
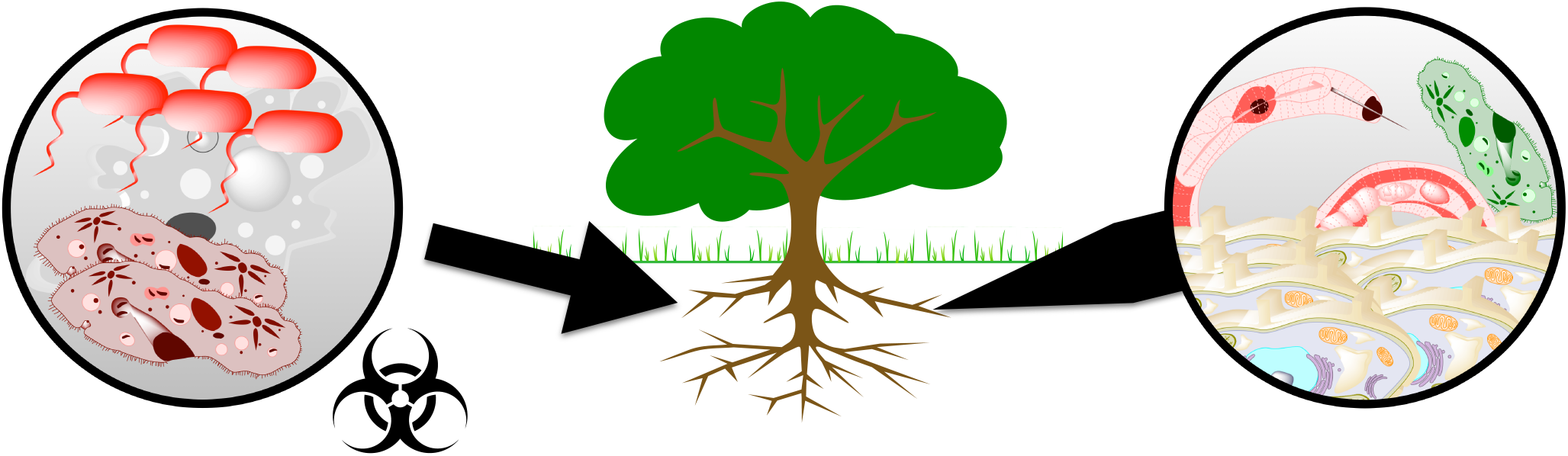
Systems approach to the Verticillium wilt of olive: a complex interaction between complex systems. Polymicrobial community attacks a host community (a host and its symbionts). Our results suggest the relevance of a systems perspective as a generalization of the approach to an infectious process.

Longitudinal (temporal) metagenomic analysis of the shotgun metagenomic sequencing data has allowed us to study the overall dynamics of the system as well as to obtain results split into amoebae and ciliates, fungi, bacteria, and nematodes. Besides, the longitudinal analysis of a root damage process has served as a real “dynamic control dataset” of the infection process afflicting the olive rhizosphere.

Our results also have very important implications in relation to the assignation of a determined parasitic species as biotroph, necrotroph or hemibiotroph. *Verticillium*, for example, has been sometimes defined as a biotrophic fungus (Boogert and Deacon 1994), whereas some other studies define it as hemibiotrophic (Zhou et al. 2012). The same happened with other fungi such as those of the *Fusarium* genus. However, our work clearly demonstrates that this kind of assignation cannot be easily done in an open, natural, and non sterile system, given the enormous complexity of an infection like the one shown in this work were both biotrophic and necrotrophic species were simultaneously intervening throughout the process. *Fusarium* is clearly recognized as hemi-biotroph simply because experiments have been conducted with plants supposedly raised from sterile seeds and plant substrates where it was assumed that only this fungus grew. In our case, we cannot get clear conclusions about it, since these plants were four month-old potted olives purchased from a commercial and non-controlled nursery (Jimenez-Ruiz et al. 2017).

Finally, this study is another example of the usefulness of the draft genomes included in the NCBI WGS database, since we have used draft genomes sequences of olive and fungi to enrich the NCBI nt database. Using the enlarged database, taxonomic classification methods correctly include the information on individual species gathered by alignment methods.

## Acknowledgments

This research received no specific grant from any funding agency in the public, commercial, or not-for-profit sectors.

## Supplemental Material

**Figure 10.**
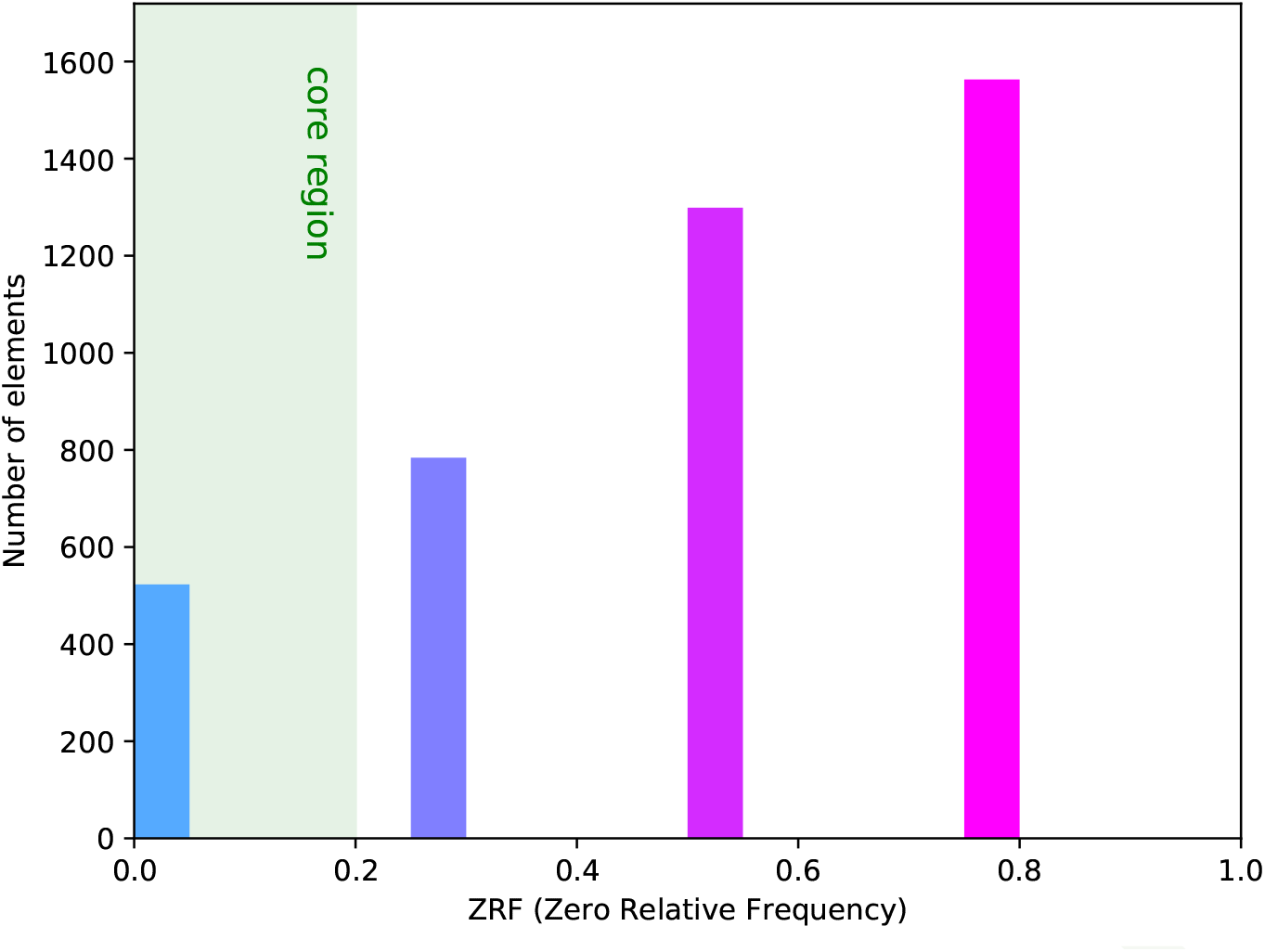
Zero relative frequency (ZRF) histogram for all the taxa of samples during the process of infection with *V. dahliae.* The ‘core’ region includes the taxa that are present in all the samples of the study, therefore, the perdurable taxa along the infection process.

**Figure 11.**
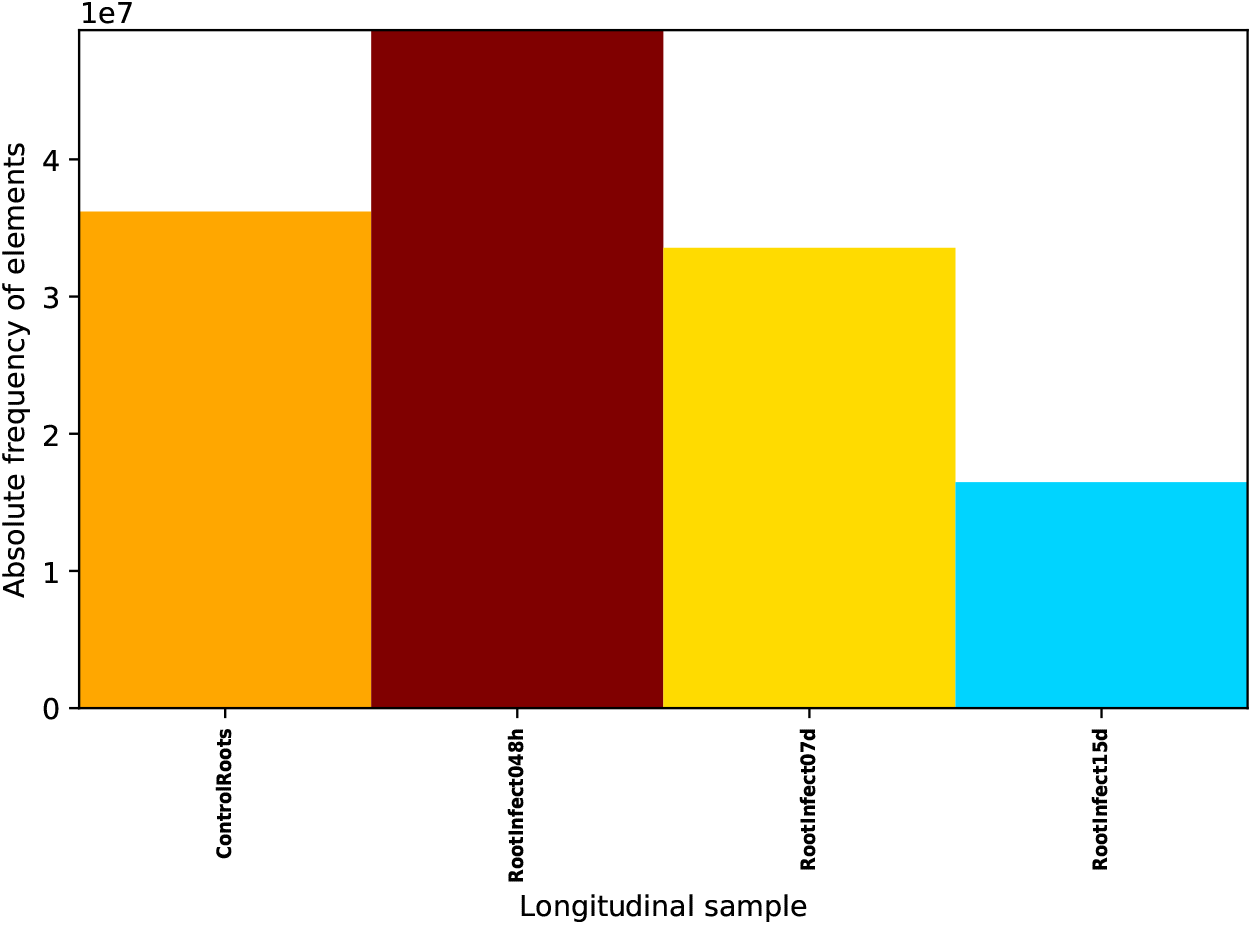
Absolute frequency plot for reads of samples during the process of infection with *V. dahliae.*

**Figure 12.**
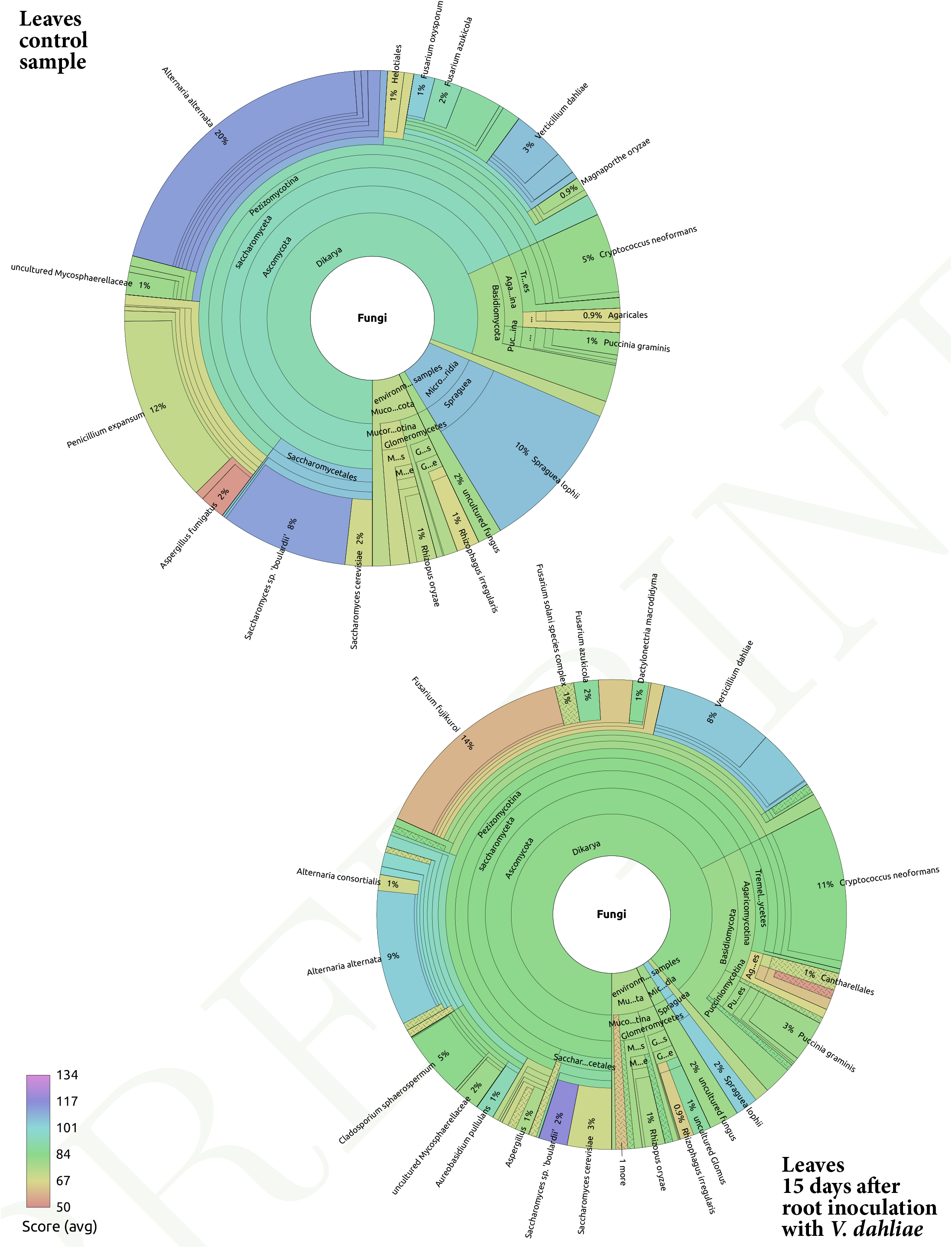
Recentrifuge plots (Marti 2018) of fungal SMS classified reads at species level for leaves during *V. dahliae* infection. The top pie corresponds to the leaves control sample while the bottom chart refers to the sample of leaves 15 days after the *Verticilium* inoculation.

**Figure 13.**
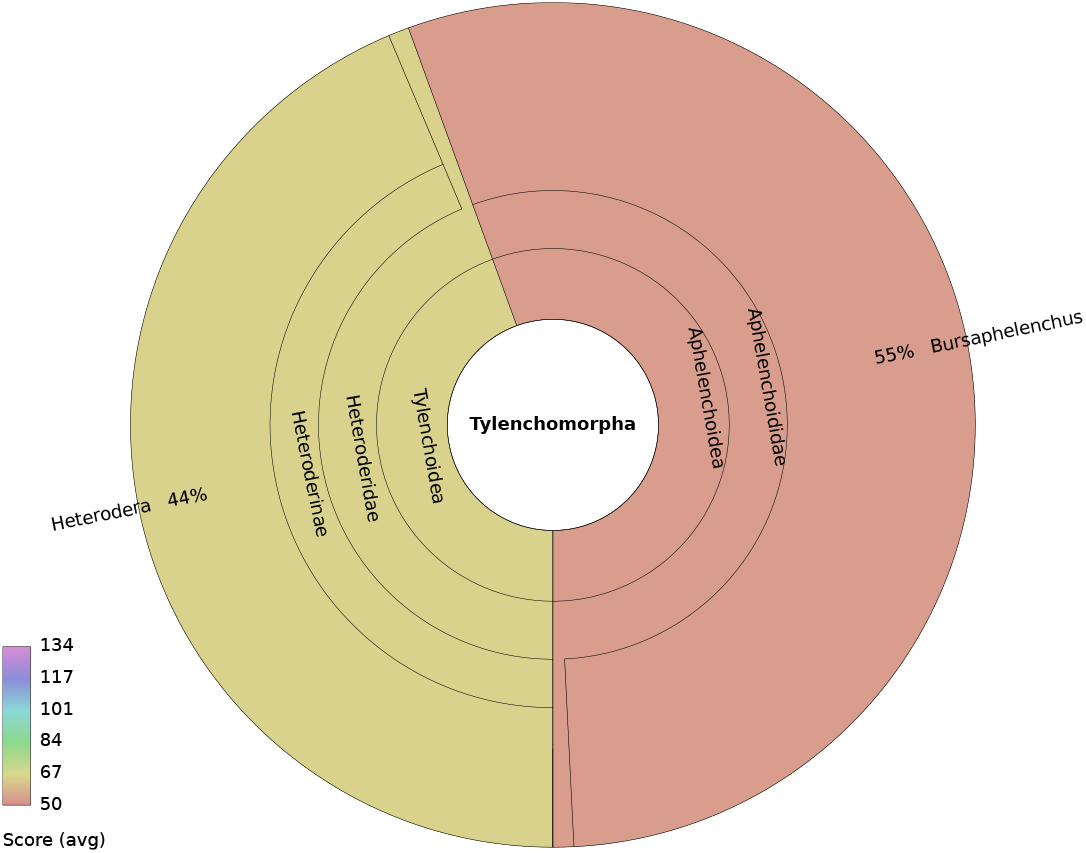
Recentrifuge plot summarizing the results for suborder Tylenchomorpha 15 days after the inoculation with *V. dahliae*. *Heretodera* and *Bursaphelenchus* genera appear with relatively low scores, but over the cutoff limit of 50.

**Figure 14.**
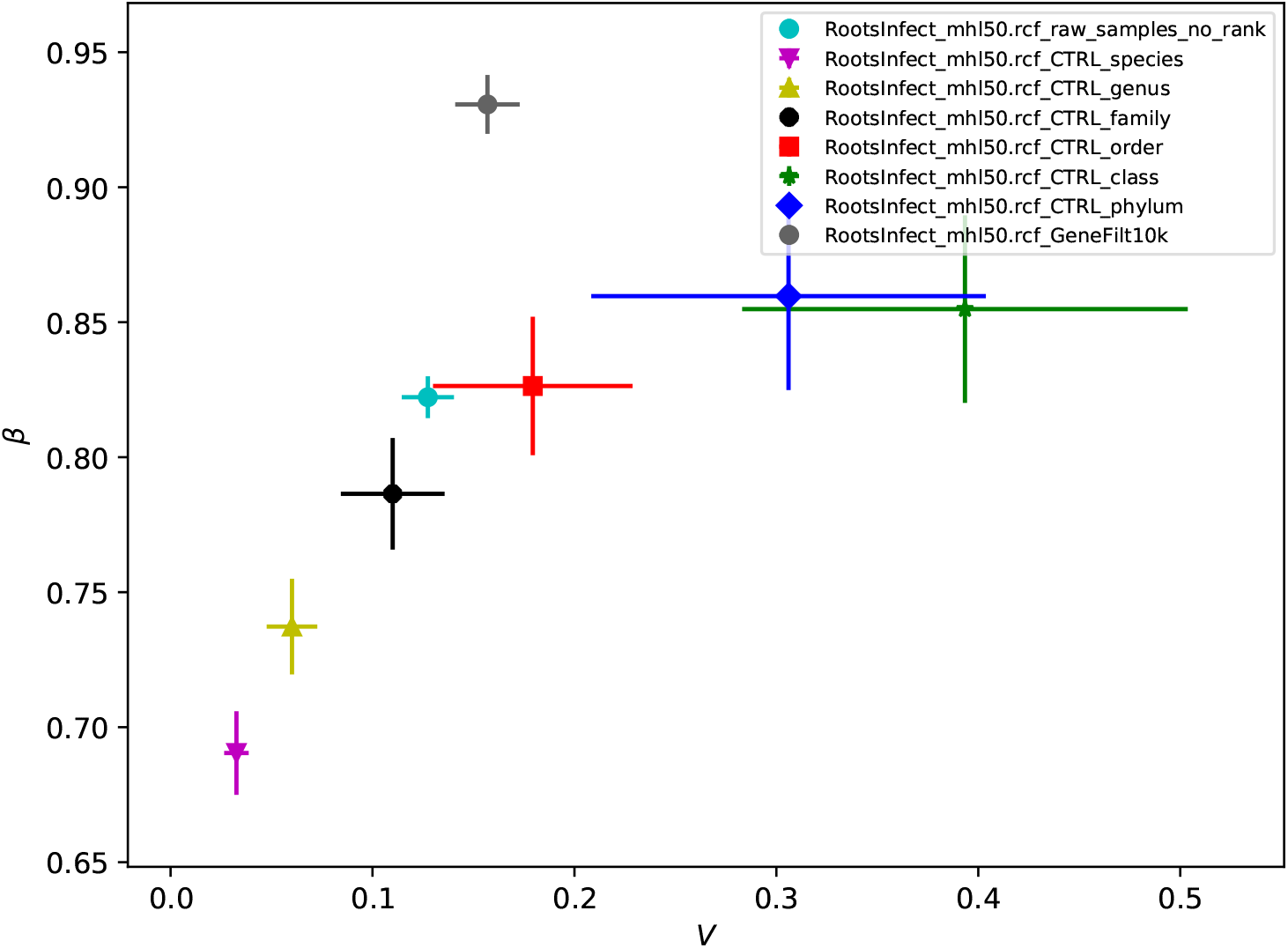
Taylor’s law parameters summary plot for x-Weighted fit during the process of infection with *V. dahliae* for various Recentrifuge datasets. See Martí et al. (2017) for details on the calculations for fitting a x-Weighted model.

**Figure 15.**
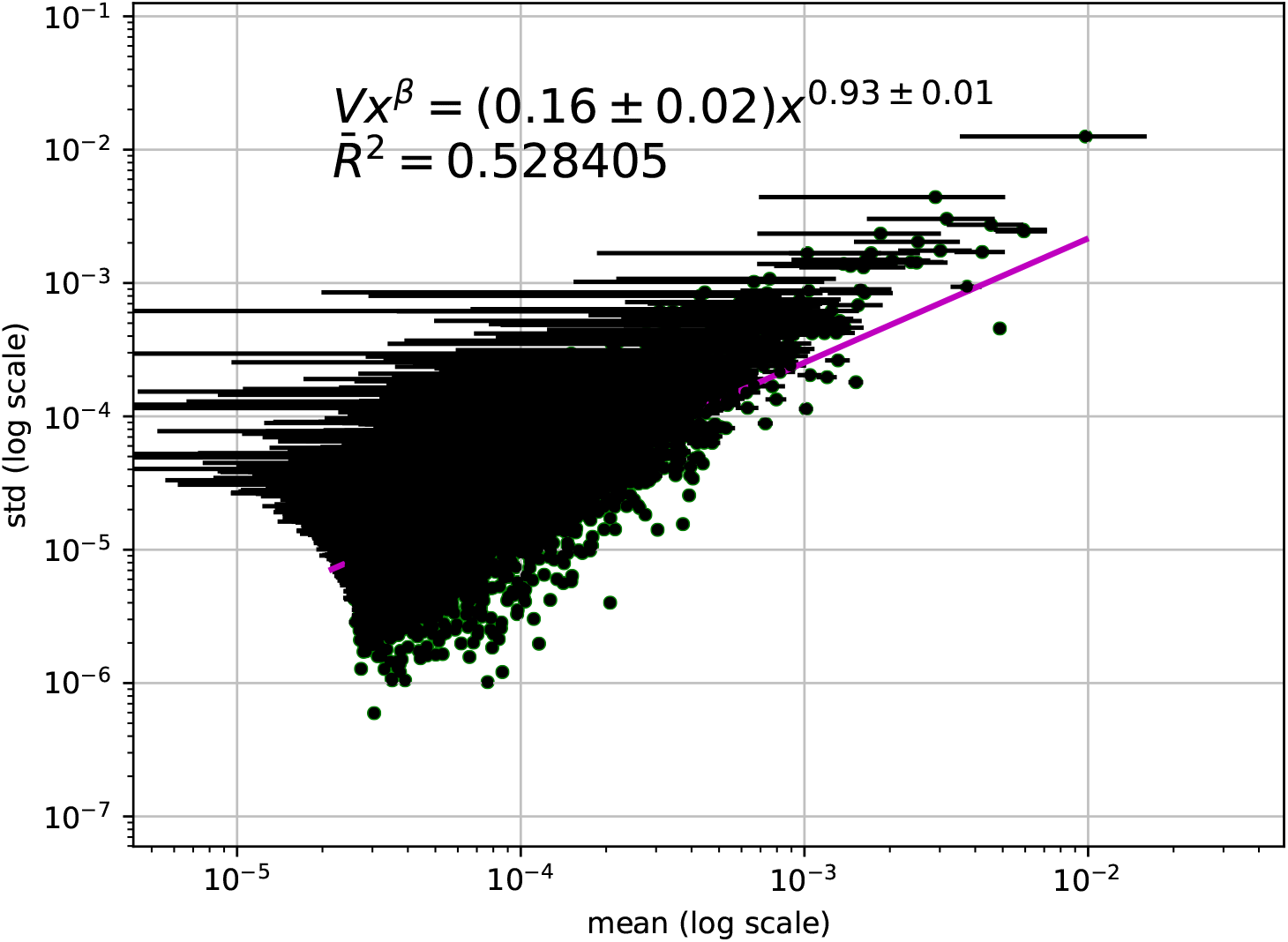
X-weighted fit for the 10,000 most representative expressed-genes during the process of infection with *V. dahliae.* See Martí et al. (2017) for details on the calculations for fitting a x-Weighted model.

**Figure 19.**
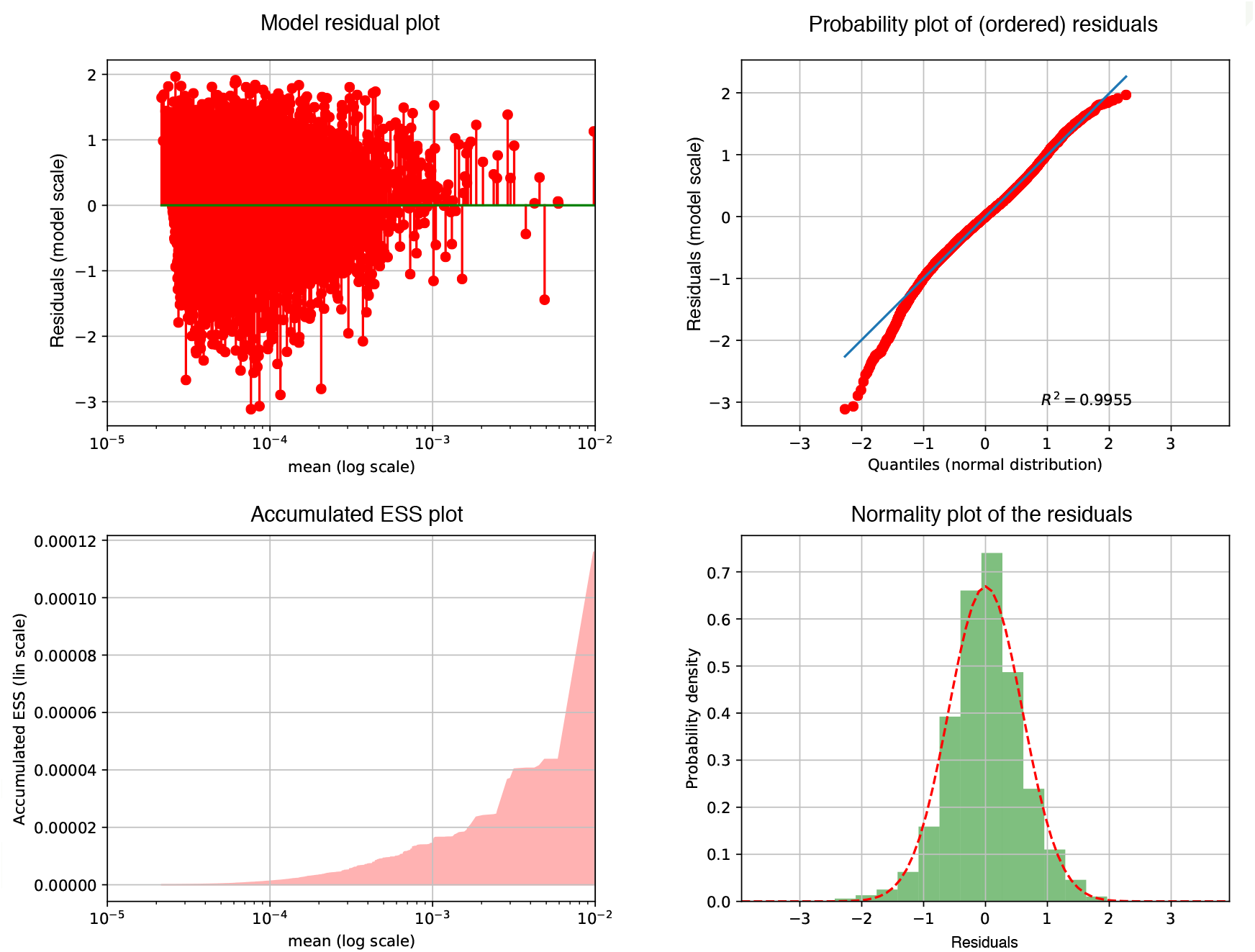
Residual analysis plot of LLR-model fit for the 10,000 most representative expressed-genes during the process of infection with *V. dahliae.* See Martí et al. (2017) for details on the calculations for fitting a x-Weighted model.

**Figure 17.**
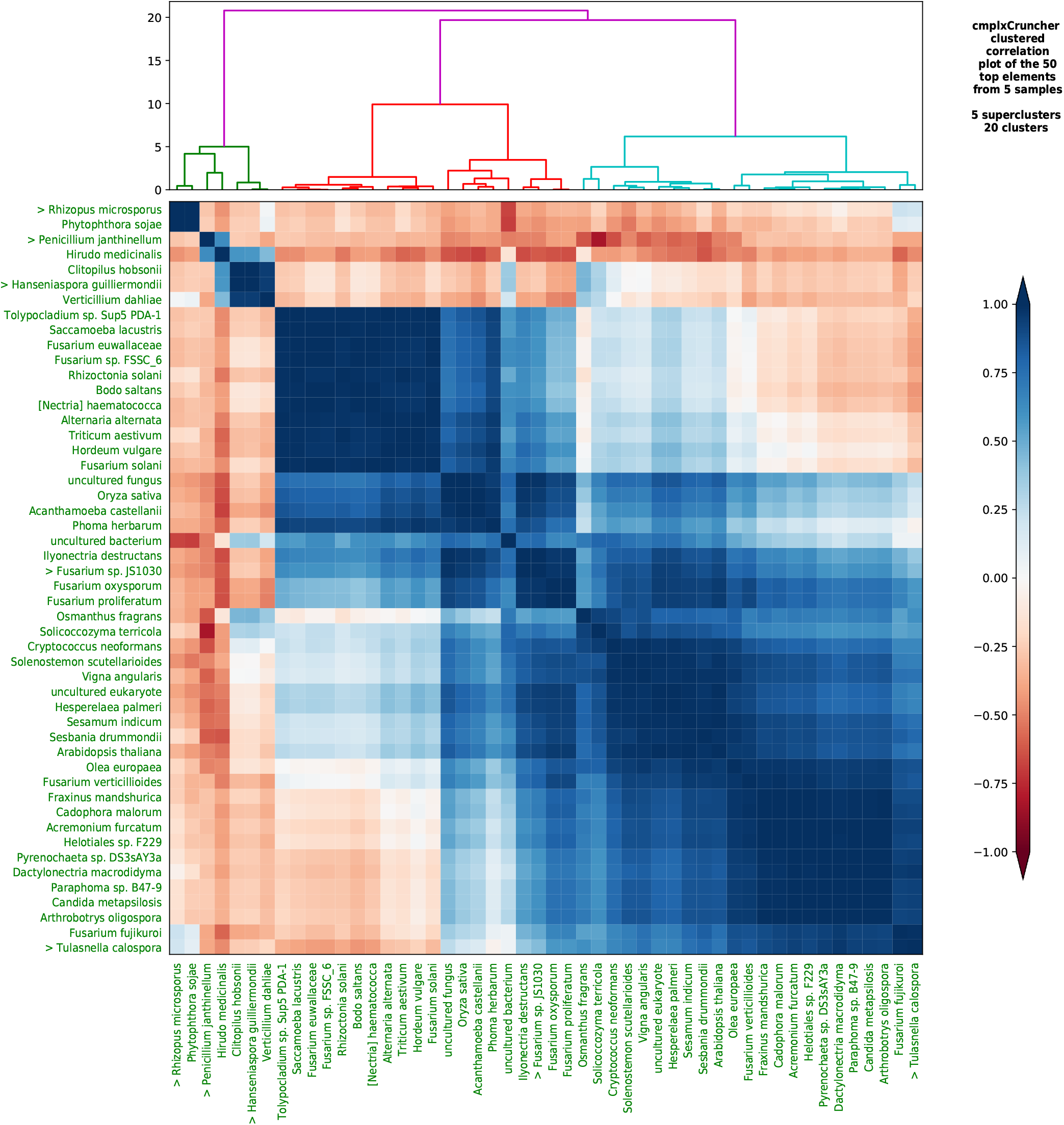
Clustered correlation and dendrogram plot for species during the process after *Olea europaea* roots damage. We show the 50 most abundant species ordered by clusterization based on the Pearson time correlation matrix. Several clusters and superclusters can be identified with this analysis by cmplxCruncher.

**Figure 18.**
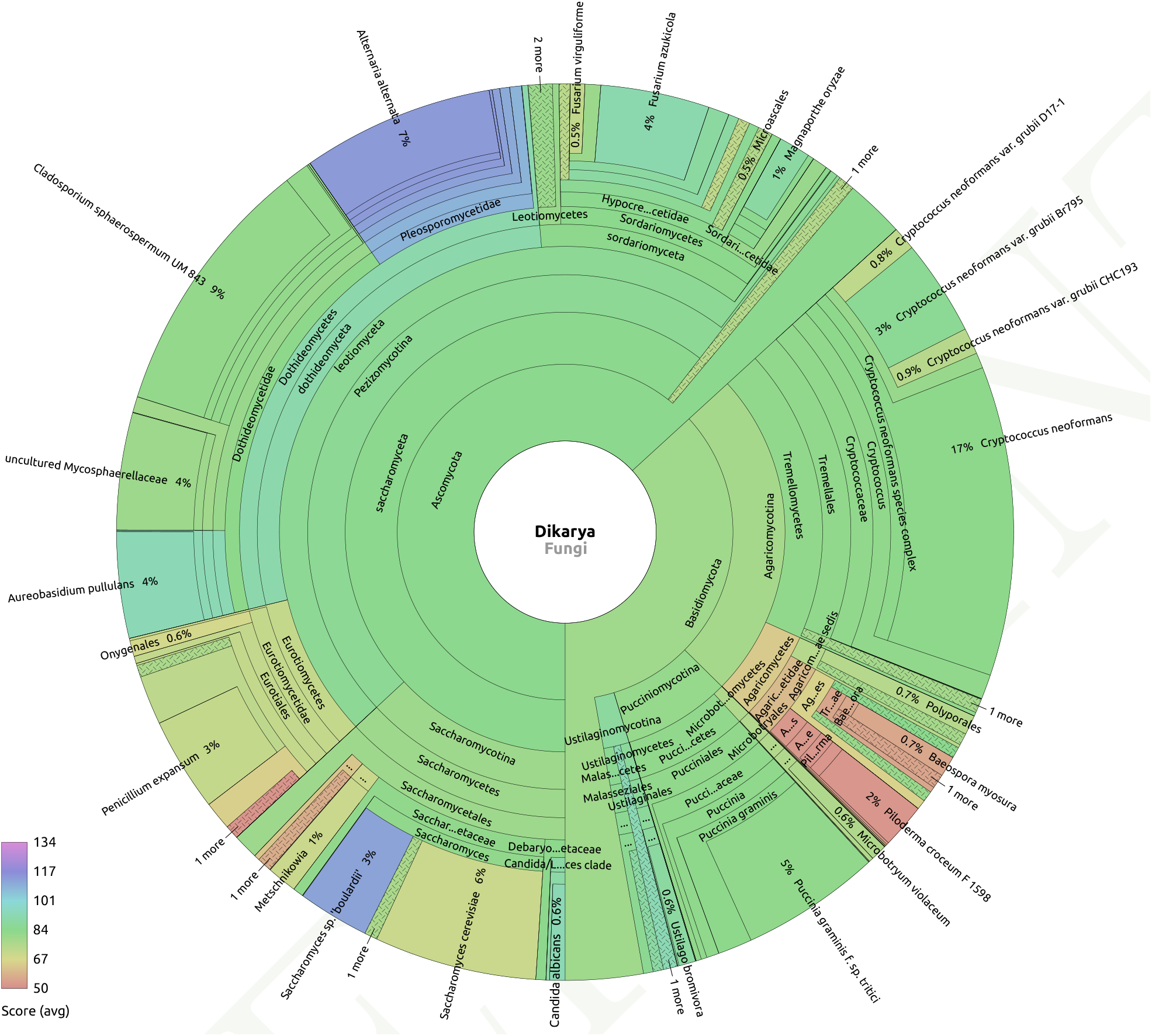
Recentrifuge plot of SMS classified reads for Dykaria fungi for the sample of leaves 15 days after the roots damage. In the bottom of the pie, *Candida albicans* appears with a good average score but low frequency.

